# Serotonin neuromodulation directs optic nerve regeneration

**DOI:** 10.1101/2024.08.12.607648

**Authors:** Kristian Saied-Santiago, Melissa Baxter, Jaffna Mathiaparanam, Michael Granato

**Author notes:** Corresponding author and lead contact: Michael Granato.

## Abstract

Optic nerve (ON) regeneration in mammalian systems is limited by an overshadowing dominance of inhibitory factors. This has severely hampered the identification of pro-regenerative pathways. Here, we take advantage of the regenerative capacity of larval zebrafish to identify pathways that promote ON regeneration. From a small molecule screen, we identified modulators of serotonin (5-HT) signaling that inhibit ON regeneration. We find several serotonin type-1 receptor genes are expressed in RGC neurons during regeneration and that inhibiting 5- HT1 receptors or components of the 5-HT pathway selectively impedes ON regeneration. We show that 5-HT1 receptor signaling is dispensable during ON development yet is critical for regenerating axons to emerge from the injury site. Blocking 5-HT receptors once ON axons have crossed the chiasm does not inhibit regeneration, suggesting a selective role for 5-HT receptor signaling early during ON regeneration. Finally, we show that agonist-mediated activation of 5-HT1 receptors leads to enhanced and ectopic axonal regrowth. Combined, our results provide evidence for mechanisms through which serotonin-dependent neuromodulation directs ON regeneration in vivo.

**Summary Statement:** 5-HT1 receptors selectively promote early optic nerve regrowth in zebrafish but are dispensable during development. Serotonin receptor-dependent modulation instructs regenerating optic nerve axons toward the brain in vivo.

## Introduction

The optic nerve is considered part of the central nervous system (CNS) and is comprised of retinal ganglion cell (RGC) axons and glial cells to relay visual information from the retina to the brain. Despite its essential role in vision, the mammalian optic nerve has limited regenerative capacity (Laha et al., 2017; Fischer et al., 2012). In response to injury or disease-induced damage, most RGCs die (Villegas-Perez et al., 1993). Even when RGCs survive the initial insult, inhibitory factors severely restrict axonal growth, resulting in loss of visual function (Silver and Miller, 2004; Buckingham et al., 2008). Combinations of growth factors have been shown to enhance the regrowth of RGC axons (Li et al., 2016; Park et al., 2008; de Lima et al., 2012); however, these axons frequently fail to properly navigate the optic chiasm and only very few reach their original brain targets (Pernet et al., 2013; Bray et al., 2017), suggesting that proper regeneration requires a more comprehensive understanding of the pathways that promote various aspects of axonal regrowth, including pathfinding and target selection. Over the past decade, it has become clear that spontaneous regeneration of injured CNS axons is not limited to non-mammalian systems but is also operant in mammals such as the naked mole rat or the spiny mouse (Park et al., 2017; Nogueira-Rodrigues et al., 2022). In these spontaneous regeneration models, optic nerve regeneration likely proceeds due to the dominance of pro-regeneration pathways, and hence, vertebrate model systems, including teleost fish, represent a unique opportunity to identify regeneration promoting pathways without the overshadowing effects of inhibitory pathways (Becker and Becker, 2007). Moreover, larval zebrafish are optically transparent, which facilitates monitoring optic nerve regeneration in vivo and in real-time (Harvey et al., 2019). Finally, larval zebrafish are amenable to small molecule screens and thus represent an unparalleled opportunity to identify factors promoting optic nerve regeneration in vivo.

Serotonin (5-HT) is a neurotransmitter that binds to 5-HT cell-surface receptors to primarily mediate communication between neurons through synaptic connections (Lipton and Kater, 1989; Zhou and Hablitz, 1999). 5-HT receptors are subdivided into seven families (Barnes and Sharp, 1999) based on their protein structure, signal transduction, and pharmacology (Hoyer et al., 1994). Recently, vertebrate studies have identified 5-HT receptors as developmental regulators of CNS axonal growth (reviewed in (Trakhtenberg and Goldberg, 2012)). For example, 5-HT receptor signaling has been shown to play instructive roles in guiding commissural axons at the CNS midline (Xing et al., 2015). In rodents, the 5-HT1B receptors and Monoamine Oxidase, an enzyme that breaks down 5-HT, are required during development to properly sort retinal projections (Upton et al., 1999; Upton et al., 2002). Finally, serotonin receptor signaling has also been implicated in post-developmental processes, including axonal regrowth. Following a spinal cord transection in adult zebrafish, 5-HT1B receptors facilitate the regeneration of spinal interneurons (Huang et al., 2021). Additionally, serotonin modulates axonal regrowth in goldfish retinal explants after an optic crush injury through the 5-HT1A and 5- HT2 receptors (Lima et al., 1994; Schmeer and Lima, 2000; Schmeer et al., 2001). Yet despite their function in modulating axonal growth and guidance, the in vivo requirement of 5-HT signaling in long-distance CNS axonal regeneration, including optic nerve regeneration, has remained elusive.

We recently developed an optic nerve transection assay in larval zebrafish that allows for live examination of the neurogenesis-independent process of spontaneous optic nerve regeneration (Harvey et al., 2019). Here, using this in vivo assay, we conducted a screen using a small molecule library against defined molecular targets (Selleckchem/Bioactive 2100). From this screen, we identified four compounds modulating serotonin signaling that impair optic nerve regeneration. Focusing on two of the identified small molecules, an agonist and an antagonist of 5-HT1 receptors, we provide compelling evidence that 5-HT1 signaling promotes axonal growth during the early stages of optic nerve regeneration. We show that blocking 5-HT1 receptors during optic nerve development does not impair developmental axon growth, arguing that 5-HT receptor signaling plays a selective role during regeneration. Lastly, we find that ectopic activation of 5-HT1 receptors leads to enhanced but also misguided regrowth of optic nerve axons, suggesting that 5-HT receptor-dependent neuromodulation plays an instructive role in directing regenerating optic nerve axons toward their original brain targets.

## Results

### A small molecule screen uncovers components of the 5-HT signaling pathway as modulators of RGC axon regeneration

To identify signaling pathways that promote RGC axon regeneration in vivo, we screened a small molecule library consisting of over 1,400 FDA approved compounds. These compounds cover a broad range of identified targets and over 40 genetic signaling pathways as defined by PANTHER pathway analysis (Mi and Thomas, 2009), including agonists and antagonists targeting all seven 5-HT receptor families and other 5-HT signaling downstream effectors. To test their in vivo roles in CNS regeneration, these compounds were applied to larvae containing the *Tg(isl2b:GFP)* transgene, which labels most, if not all, RGC neurons and their axons express GFP (Figures 1A-B)(Pittman et al., 2008). Injury was induced when larvae were 5-day-old post-fertilization (5 dpf), a time point when their visual system is already functional (Brockerhoff et al., 1995; Easter and Nicola, 1996). In these animals, we fully transected the optic nerve halfway between the retinal exit point and the optic chiasm using a sharpened tungsten needle. We previously showed that regrowing RGC axons innervate both ipsilateral and contralateral tecta following complete axonal transection (Harvey et al., 2019). Therefore, to simplify analysis and ensure the regenerating axons from a single injured nerve were scored properly, we removed the uninjured right eye (Figure 1B’’). After transection, the transected optic nerve undergoes a stereotypic timeline of recovery. By 24 hours post-transection (hpt), the nerve segment distal to the injury site undergoes fragmentation/ degeneration, and regenerating optic nerve axons emerge from the nerve stump and begin to extend toward the optic chiasm located at the CNS midline (Figure 1C)(Harvey et al., 2019). By 48 hpt, regrowing RGC axons in wild-type larvae have extended past the optic chiasm and have begun to re-innervate the peripheral edges of the optic tecta (Figure 1D-D’). In order to assess the in vivo effects of the small molecule library, particularly on early axon guidance and extension, we applied small molecules (or a DMSO control) to larvae 24 hpt after full transection of the optic nerve (Figure 1C’). From the small molecule library, we identified and independently confirmed four small molecules known to modulate serotonin signaling that resulted in impaired optic nerve regeneration. One identified molecule is the endogenous ligand serotonin, while another, tranylcypromine, is an antagonist that inhibits the monoamine oxidase and the lysine-specific demethylase 1 (LSD1) enzymes (Table 1). The remaining two molecules represent selective agonists and antagonists against 5-HT1 receptors. Specifically, WAY-100635 is a 5-HT antagonist that inhibits type-1A receptors, while zolmitriptan has been validated as an agonist that activates 5-HT type-1B/1D receptors (Table 1). Given this selectivity, we focused on further determining the roles of 5-HT1 receptors in optic nerve regeneration.

**Figure 1.**
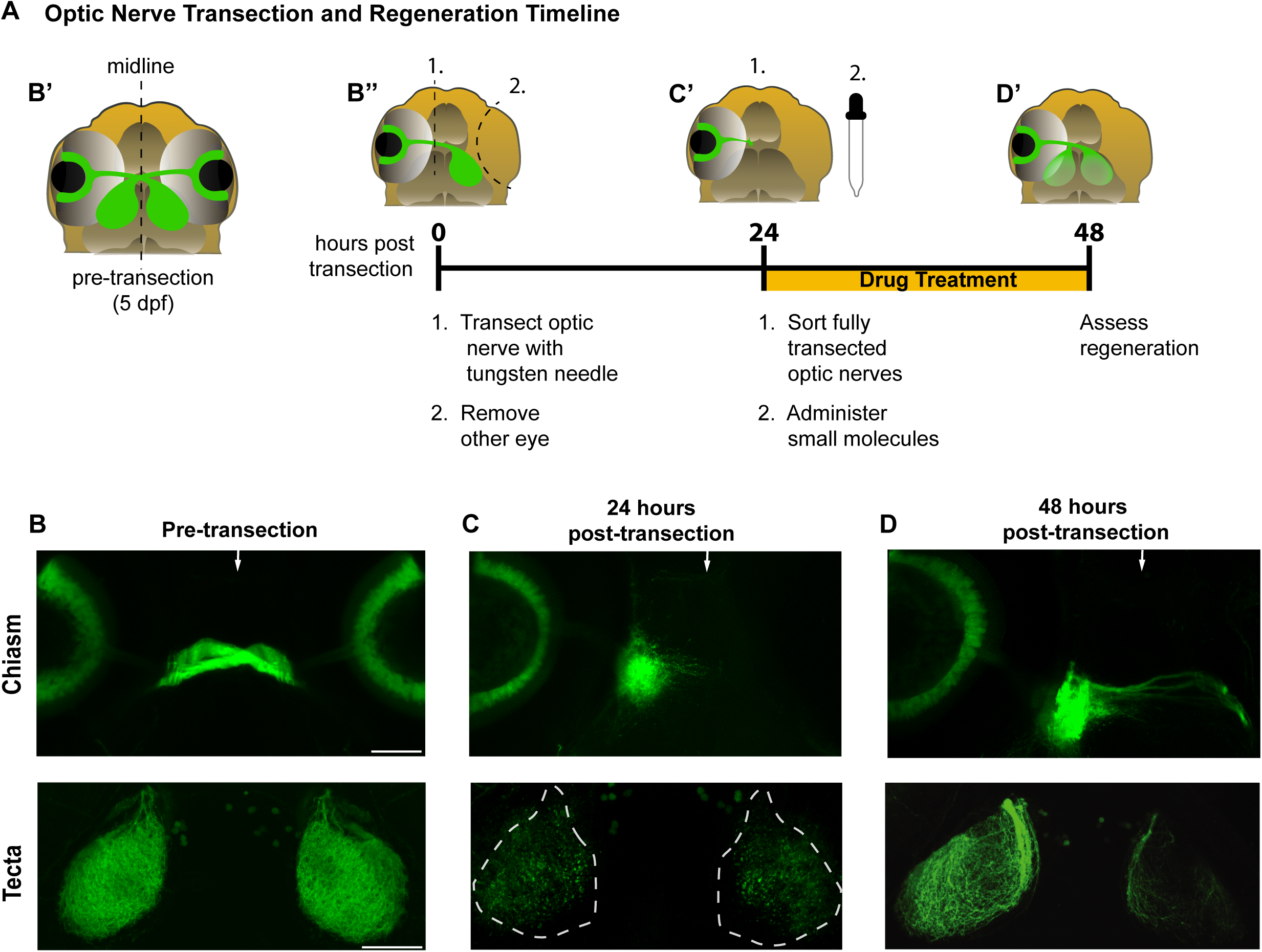
Regenerating RGC axons robustly regrow to the optic tecta by 48 hours post-transection. (A) Timeline of optic nerve transection, regeneration, and larvae exposure to small molecules. Schematic describes the pre-transection and post-transection (0, 24 & 48 hpt) time points during optic nerve regeneration and its assessment in larval zebrafish. (B - D) Fluorescent representative images of *Tg(Isl2b:GFP)* fixed larvae visualized using confocal imaging at distinct time points: pre-transection, 24 hrs post-transection, and 48 hrs post-transection. Arrows depict the midline at the optic chiasm. Different larvae were used across time points. Scale bars 50 uM. (B-B’) At 5 days post-fertilization (5 dpf), RGC axons have crossed the midline at the optic chiasm (top panel) and innervated the contralateral tecta (bottom panel). (B”) At 0 hpt, the left optic nerve of *Tg(Isl2b:GFP)* larvae were transected and the right eye was removed as detailed in Material and Methods. (C-C’) At 24 hpt, transected axons begin to extend towards the optic chiasm (chiasm region, top panel). Full transection of the optic nerve was corroborated by examining larvae dorsally (tecta region, bottom panel) at 22-24 hpt for remnants of Isl2b:GFP nerve structures. Degenerated tecta are outlined with dashed lines. (D-D’) At 48 hpt, wild-type larvae regrow their RGC axons toward the optic chiasm (top panel) and begin re-innervating the peripheral edges of the tecta (bottom panel). Live zebrafish larvae treated with small molecules were assessed to determine whether optic nerve regeneration was impaired. Top and bottom panels for each time point are of the same larva. At this time point, the ipsilateral tectum (left tectum) is typically more innervated than the contralateral tectum.

**Table 1:** Small molecules targeting the 5-HT signaling pathway impair optic nerve regeneration.

| Small Molecule | Type of Small Molecule | Target(s) | Phenotype |
| --- | --- | --- | --- |
| WAY-100635 | Antagonist | 5-HT1A receptor | Reduced tectal re-innervation |
| Zolmitriptan | Agonist | 5-HT1B/1D receptor | Increased ectopic regrowth |
| Tranylcypromine | Antagonist | Monoamine oxidase/<br>LSD1 | Reduced tectal re-innervation |
| Serotonin | Agonist | Multiple 5-HT receptors | Reduced tectal re-innervation |

### Inhibiting 5-HT1 receptors significantly reduces RGC regenerative axonal growth

We first characterized the optic nerve regeneration phenotype observed after treatment with the small molecule WAY-100635. This serotonin receptor antagonist is a well characterized inhibitor of 5-HT1A receptors in vitro (Ferreira et al., 2010) and in vivo (Maximino et al., 2013; Long et al., 2023), with 100-fold higher binding selectivity to 1A receptors over other subtypes (Forster et al., 1995). We exposed larvae with fully transected optic nerves to vehicle (0.3% DMSO) or WAY-100635 from 24 - 48 hpt, and at 48 hpt quantified the number of transected optic nerves that had extended to the contralateral tectum. Compared to DMSO-treated control animals, we observed a significant and dose-dependent reduction of optic nerve regrowth in WAY-100635 treated larvae (compare Figures 2A-D, DMSO controls with 5 uM or 50 uM WAY-100635 treated larvae). Specifically, after incubating larvae in 50 uM WAY-100635 media, we observed a 2.2-fold increase of nerves (Figure 2D) that stalled at various positions along their regenerative path (Figures 2B-C, yellow arrowheads in chiasm panels), and failed to re- innervate the contralateral tectum (Figures 2B-C, white dashed lines in tecta panels). To determine whether RGC axons in animals treated with 50 uM WAY-100635 stall before or after the optic chiasm, we examined whether optic nerves crossed the optic chiasm by 48 hpt. When compared to control animals, regenerating optic nerves in WAY-100635 treated larvae failed to cross the optic chiasm at a significantly higher rate (42% in 50 uM WAY-100635 treated larvae compared to 19% in control nerves (Figure S1A, black bars)). Lastly, to determine whether the growth defects observed in optic nerves of WAY-100635 treated animals are due to regenerating axons deviating from their pre-transection trajectory or stalling prematurely along their trajectory path, we quantified the number of optic nerves displaying ectopic axonal regrowth (Figure 2B, yellow arrow). After treatment with low (5 uM) or high (50 uM) concentration of the 5-HT1 antagonist WAY-100635, we failed to observe a significant increase in ectopic axonal regrowth (Figure 2E), consistent with the idea that WAY-100635 causes RGC axons to stall before or at the optic chiasm. Combined these results strongly suggest that 5-HT1 receptors promote RGC axonal regrowth.

**Figure 2.**
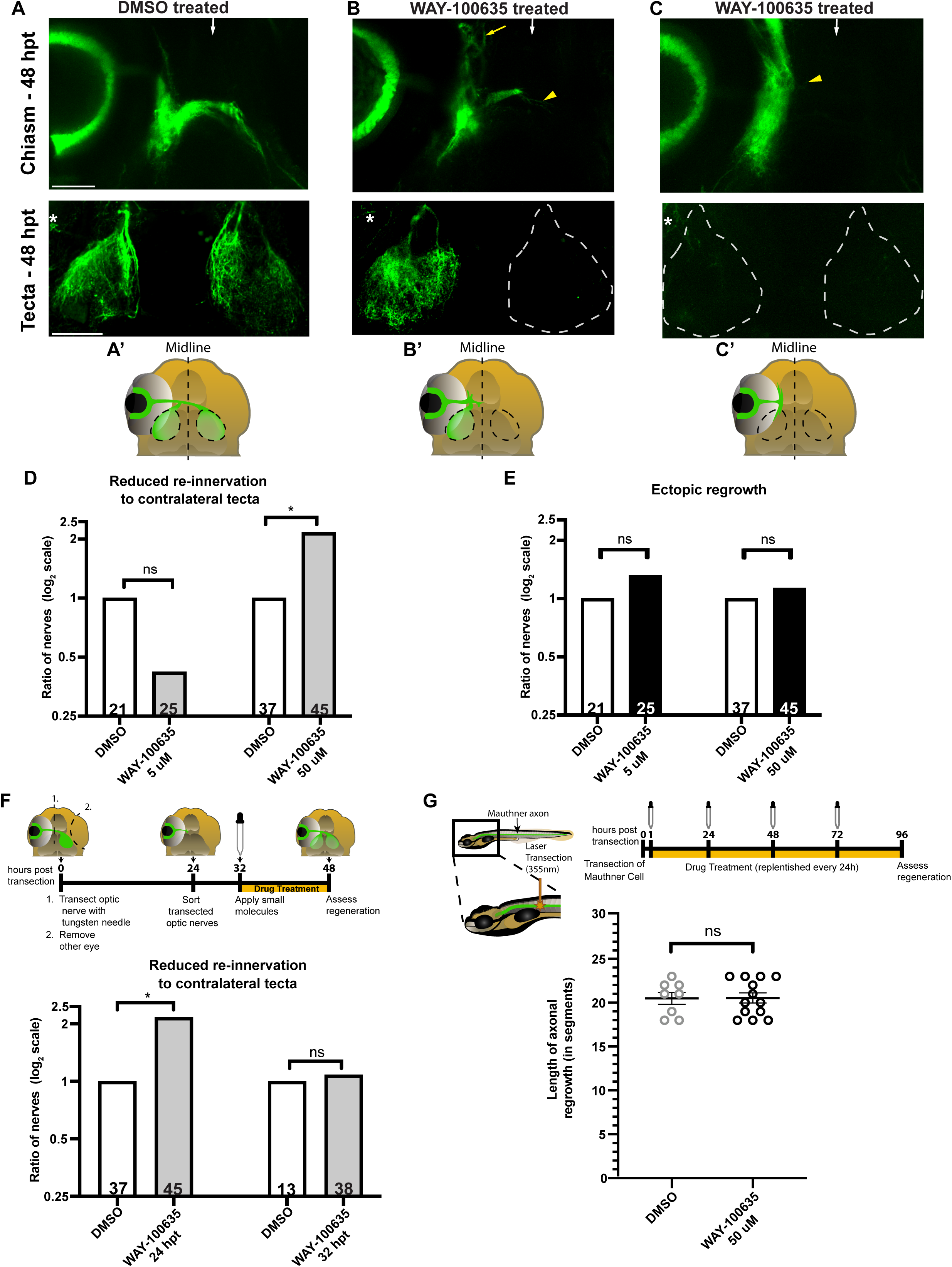
5-HT1 receptor signaling promotes early optic nerve regrowth without impairing overall CNS axonal regrowth. (A - C) Fluorescent representative images and (A’-C’) schematics of *Tg(Isl2b:GFP)* at 48 hpt of larvae exposed to (A) DMSO 0.3% and (B & C) 50 uM of antagonist WAY-100635. Images in B and C are examples of two different types of regrowth observed when applying this small molecule. White arrows depict the midline region at the optic chiasm. Asterisks depict Isl2b:GFP staining outside of the optic tecta. Scale bars = 50 uM. (B) RGC axons of larvae treated with the antagonist WAY-100635 regrew toward the optic chiasm (top panel, yellow arrowhead) but failed to innervate the contralateral tectum (dashed lines, bottom panel). A group of axons regrew ectopically away from the optic chiasm (yellow arrow). Re-innervation of the ipsilateral tectum begins soon after the small molecules are added. (C) Regenerating RGC axons of a larva treated with the antagonist WAY-100635 stalled near the injury site (yellow arrowhead, top panel). No axonal regrowth is observed to the optic chiasm or contralateral tectum (dashed lines, bottom panel). (D) Quantification of optic nerve axonal re-innervation to contralateral tectum at 48 hpt in *Tg(Isl2b:GFP)* larvae treated at 24 hpt with DMSO 0.3% or WAY-100635. Bars represent the ratio of optic nerves that fail to re-innervate the contralateral tectum in WAY-100635 treated (gray bars) and control groups, plotted as a log2 scale. See Material and Methods for details on ratios and fold-change calculations. Statistics determined using the two-tailed Fisher’s exact test. Asterisks in figures represent statistical significance: * *P* < 0.05; ns, not significant. n=21 and n=25 for nerves treated with DMSO and 5 uM WAY-100635, respectively; n=37 and n=45 for nerves treated with DMSO and 50 uM WAY-100635, respectively. (E) Quantification of ectopic regrowth at 48 hpt in *Tg(Isl2b:GFP)* larvae treated at 24 hpt with DMSO 0.3% or WAY-100635. Bars represent the ratio of optic nerves showing ectopic regrowth in WAY-100635 treated (black bars) and control groups, plotted as a log2 scale. See Material and Methods for details on ratios and fold-change calculations. Data displayed in Figure 2D-E was obtained from larvae in the same treated groups. Statistics determined using the two-tailed Fisher’s exact test. ns, not significant. n=21 and n=25 for nerves treated with DMSO and 5 uM WAY-100635, respectively; n=37 and n=45 for nerves treated with DMSO and 50 uM WAY-100635, respectively. (F) (Top panel) Timeline of optic nerve regeneration from 0 hpt - 48 hpt. DMSO or small molecules were added to *Tg(Isl2b:GFP)* larvae at 32 hpt and optic nerve regeneration was assessed at 48 hpt. Larvae were kept in 1x PTU/E3 from 24 hpt - 32 hpt. (Bottom panel) Quantification of optic nerve axonal re-innervation to contralateral tecta at 48 hpt in *Tg(Isl2b:GFP)* larvae treated with DMSO 0.3% or 50 uM WAY-100635 at 24 and 32 hpt (gray bars). Bar graphs and the ratios observed were calculated as detailed in Figure 2D and Material and Methods. Data showing 50 uM WAY-100635 at 24 hpt is identical to Figure 2D and shown here for visual comparison only. Statistics determined using the two-tailed Fisher’s exact test. Asterisks in figures represent statistical significance: * *P* < 0.05; ns, not significant. n=37 and n=45, for nerves treated with DMSO and WAY-100635 at 24 hpt, respectively. n=13 and n=38, for nerves treated with DMSO and WAY-100635 at 32 hpt, respectively. (G) (Top panel) Timeline of mauthner axon regeneration. Schematic shows a laser transection (orange laser) performed in *Tg(Tol-056:GFP)* 5 dpf larvae using an UV-laser (355-nm wavelength) at the ninth spinal cord hemisegment. After transection, DMSO 0.3% or 50 uM WAY-100635 was added to larvae starting at 1 hpt (dropper). Fresh DMSO and small molecules were replenished at 24, 48 and 72 hpt. Mauthner axon regrowth was assessed at 96 hpt. (Bottom panel) Quantification of axonal length regrowth at 96 hpt, measured in spinal cord segments in control and WAY-100635 treated groups. Statistics determined using the one-way ANOVA test. ns, not significant.

To determine whether 5-HT1 receptors promote RGC axonal regrowth along the entire regenerative trajectory to the optic tectum or selectively at distinct portions of the regenerative path, we blocked 5-HT1 receptors once RGC axons had reached the optic chiasm. For this, we applied the antagonist WAY-100635 to larvae with fully transected optic nerves at 32 hpt, and assessed contralateral tectal re-innervation at 48 hpt (Figure 2F, top panel). We found that optic nerves in larvae treated with 50 uM WAY-100635 re-innervated the contralateral tectum to a similar extent as that observed in DMSO controls (Figure 2F, bottom panel). Together with our findings that inhibition of 5-HT1 receptors at 24 hpt significantly reduces the re-innervation of RGC axons to the contralateral tecta (Figure 2D), these results provide compelling evidence that 5-HT1 receptors promote RGC axon regrowth during the early stages of axon regeneration once axons emerge from the nerve stump and extend towards the optic chiasm.

We next examined whether 5-HT1 receptor signaling acts to promote axonal regrowth broadly within the CNS or whether it selectively promotes optic nerve regeneration. For this, we examined the role of 5-HT1 receptors in spinal cord regeneration. Specifically, we focused on a pair of reticulospinal neurons, the Mauthner cell neurons, to quantitatively and at single-neuron resolution determine the effects of 5-HT1 receptor antagonists in spinal cord regeneration. A single mauthner cell neuron resides on either side of the hindbrain, and its axon extends from the hindbrain to the tip of the spinal cord (Eaton et al., 1977). We used a well-established assay to laser-transect mauthner axons in 5 dpf larvae (Bremer et al., 2019) to test whether blocking 5-HT1 receptors impairs mauthner axonal regeneration. We added 50 uM WAY-100635 at 1 hpt and then replaced the WAY-100635 containing incubation media every 24 hours until 72 hpt (Figure 2G, top panel). We found that compared to DMSO-treated control animals, the length of mauthner axonal regrowth at 96 hpt in larvae treated with the 5-HT1 antagonist WAY-100635 was indistinguishable (Figure 2G, bottom panel), demonstrating that Mauthner cell axon regeneration is unaffected by the presence of 5-HT1 receptor inhibitors. Combined with the effect of 5-HT1 receptor signaling on RGC regeneration, our results suggest a selective role for 5-HT1 receptors in RGC axon regeneration.

### 5-HT1 receptor genes are expressed in RGC neurons during RGC axon regeneration

5-HT1 receptors are expressed in the vertebrate CNS, including in the retina of mice and zebrafish larvae (Upton et al., 1999; Norton et al., 2008). Given the apparent selective role of 5- HT1 receptors in RGC axonal regeneration, we wondered whether 5-HT1 receptors are expressed in RGC neurons during regeneration. To detect mRNA expression of 5-HT1 receptors in the RGC layer, we performed whole-mount fluorescent *in situ* hybridization (FISH) with a hybridization chain reaction (Choi et al., 2018). We obtained specific probes for the four zebrafish 5-HT1 receptors targeted by small molecules that impaired optic nerve regeneration from our screen: *htr1aa, htr1ab, htr1b,* and *htr1d.* After incubating *Tg(isl2b:GFP)* 5 dpf larvae with a probe mix against all four *htr1* receptors prior to optic nerve transection, we detected expression of these serotonin receptors distributed across the RGC layer (compare Figures 3A- C, no probe with Figures 3D-F, four *htr1* receptors) and in Isl2b:GFP RGC neurons (Figure 3F’, yellow dashed circles). Next, we incubated uninjured larvae or larvae that were fixed 24 hrs after transecting both of their optic nerves with a probe mix against *htr1aa* and *htr1ab,* receptors targeted by the antagonist WAY-100635. We detected expression of these receptors in RGC neurons before transection (Figures 3G-I and yellow dashed circles in Figure 3I’, uninjured) and also during regeneration at 24 hpt (Figures 3J-L and yellow dashed circles in Figure 3L’, transected). Thus, 5-HT1A receptor genes are expressed in RGC neurons at a critical time when 5-HT1 receptors promote RGC axonal regrowth towards the optic chiasm.

**Figure 3.**
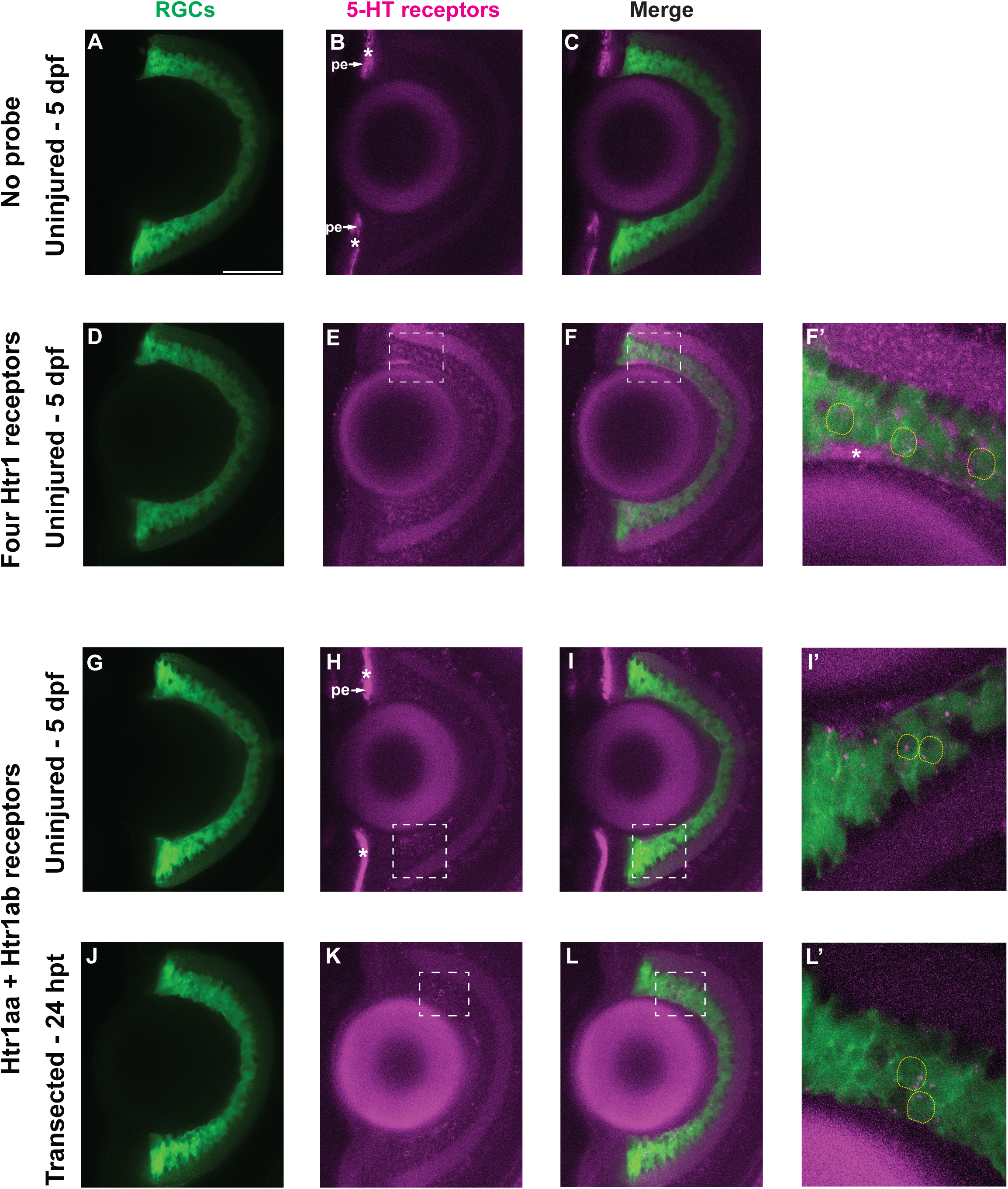
5-HT1 receptor genes are expressed in RGCs at pre-transection and during optic nerve regeneration. (A - F) Representative images of retinas from *Tg(isl2b:GFP)* 5 dpf uninjured larvae stained with no probe control (A-C, n=8 retinas) or an *in situ* hybridization HCR probe mix for *htr1aa, htr1ab, htr1b* and *htr1d* (D-F’, n=12)(magenta). Images shown are a maximum Z-projection of 6 optical sections (32 um), 40X. Dashed white boxes in (Figures E-F) is the area enlarged 1.5X in (F’), a merged maximum Z-projection of 10 optical sections (0.1 um) showing multiple *Tg(isl2b:GFP)* RGC neurons (green, outlined with yellow dashed line) expressing mRNA of 5-HT1 receptors (magenta). In (F’) the brightness of the green channel was adjusted for better visualization of 5- HT1 receptor genes inside RGC neurons. White asterisks depict unspecific staining. The retinal pigmented epithelium (pe) was nonspecifically labeled by the amplifier with Alexa Fluor 546. Scale bars = 50 uM. (G - L) Representative images of retinas from *Tg(isl2b:GFP)* 5 dpf uninjured larvae (G-I’, n=7) or larvae with transected optic nerves (J-L’, n=5) at 24 hpt stained with an *in situ* hybridization HCR probe mix for *htr1aa* and *htr1ab* (magenta). Images shown are a maximum Z-projection of 7 optical sections (G-I) and 12 optical sections (J-L)(32 um), 40X. Dashed white boxes in (Figures H-I) is the area enlarged 1.5X in (I’), a merged maximum Z-projection of 12 optical sections (0.1 um). In (I’), yellow dashed lines outline cell bodies with mRNA expression. The brightness of the green channel was adjusted for better visualization of 5-HT1 receptor genes inside RGC neurons. Dashed white boxes in (Figures K-L) is the area enlarged 1.5X in (L’), a merged maximum Z-projection of 11 optical sections (0.1 um). In (L’), yellow dashed lines outline cell bodies with mRNA expression. The brightness of the green channel was adjusted for better visualization of 5-HT1 receptor genes inside RGC neurons. White asterisks depict unspecific staining. The retinal pigmented epithelium (pe) was nonspecifically labeled by the amplifier with Alexa Fluor 546.

### 5-HT1 receptor signaling is dispensable for developmental RGC axonal growth

Differentiation of RGC neurons begins around 28 hours post-fertilization (hpf), and soon thereafter, RGC axons exit from the retina and extend to the optic chiasm (Laessing and Stuermer, 1996). By 48 hpf, the optic fissure closes, and most RGC axons have arrived at the optic tectum (Figure 4A)(Stuermer 1988). Expression of 5-HT1 receptor genes has previously been detected in the developing retina and tectum of zebrafish (Pei et al., 2016). Since 5-HT1 receptors are expressed in RGC neurons in 5 dpf larvae, we asked whether 5-HT1 receptor-dependent signaling is also critical to promote RGC axonal growth during development. Considering the developmental timeline of RGCs, we began treating embryos with the antagonist WAY-100635 at 24 hpf. We then examined whether developing optic nerves had reached the optic tecta at 48 hpf (Figure 4A). The innervation of optic nerve axons in WAY-100635 treated embryos was indistinguishable when compared to DMSO-treated controls (Figures 4B-D). Therefore, we conclude that 5-HT1 receptor function is likely dispensable in development for RGC axons to project to the optic tectum.

**Figure 4.**
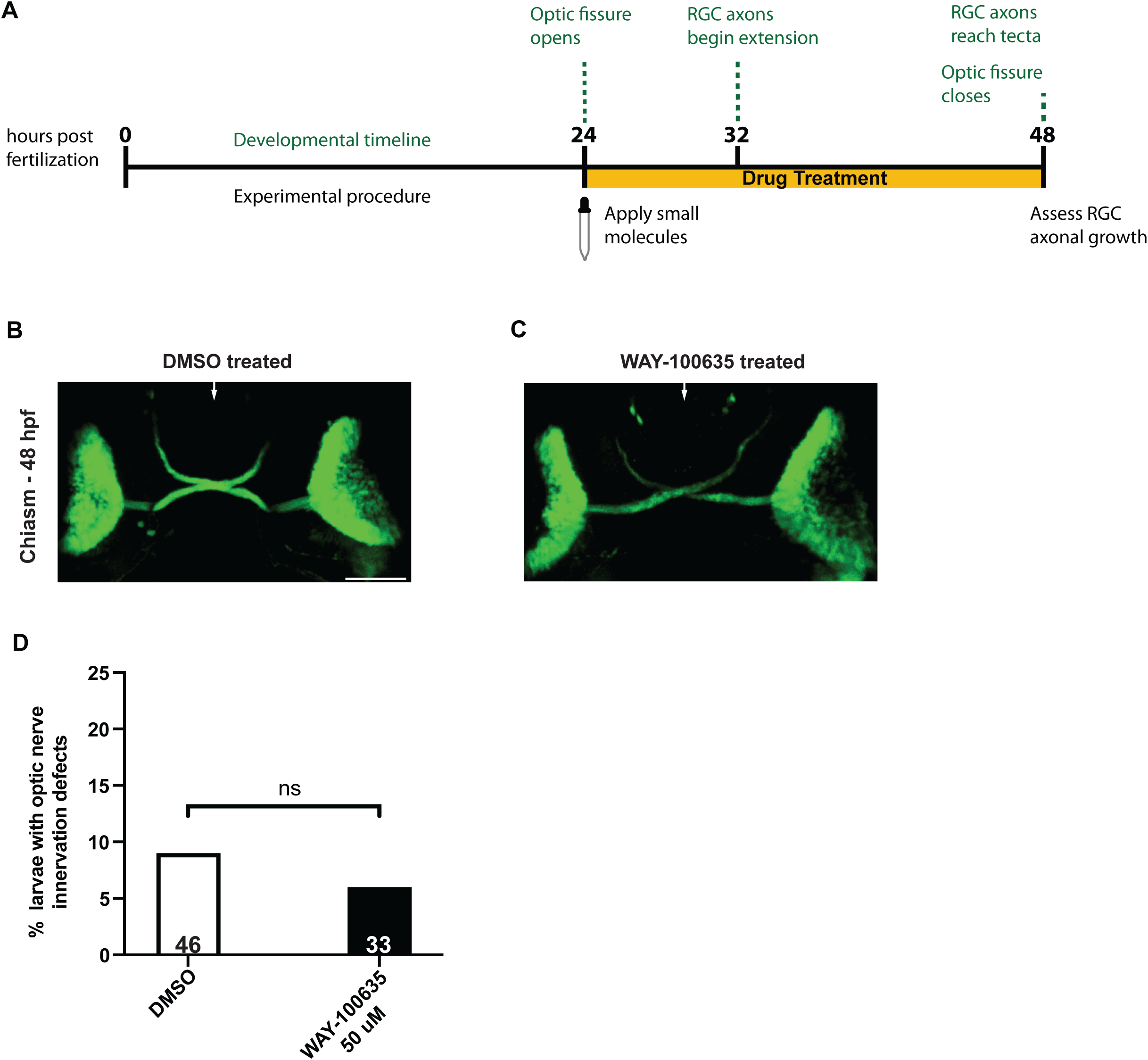
Optic nerve development is unaffected after treating zebrafish embryos with 5- HT1 antagonists. (A) Timeline of optic nerve development from 24 - 48 hours post-fertilization (hpf) and larvae exposure to small molecules. DMSO 0.3% or 50 uM of antagonist WAY-100635 was added to embryos at 24 hpf. Around 32 hpf, RGCs differentiate and immediately begin extending towards the brain. At 48 hpf, RGC axons have extended past the optic chiasm and have begun innervating the optic tecta. At this time point, treatment was washed off and optic nerve development was assessed. Timeline adapted from (Chhetri et al., 2014). (B - C) Representative images of 48 hpf *Tg(Isl2b:GFP)* embryos treated with DMSO 0.3% (B) and 50 uM WAY-100635 (C). White arrows depict the midline at the optic chiasm. No apparent growth or guidance defects are observed in the developing optic nerves of control and small molecule treated larvae. Scale bars = 50 uM. (D) Quantification of optic nerve growth defects at 48 hpf in *Tg(Isl2b:GFP)* larvae treated with DMSO or WAY-100635 (black bar). Bars represent the percentage of defective optic nerves for each treatment group. Statistics determined using two-tailed Fisher’s exact test. Statistical significance: ns, not significant. n=46 and n=33, for larvae treated with DMSO and WAY-100635, respectively.

### 5-HT1 receptors direct regenerating RGC axons towards the optic chiasm

We next asked whether 5-HT1 receptor signaling plays an instructive role during optic nerve regeneration. To test this, we used the 5-HT1B/1D selective agonist Zolmitriptan (Wurch et al., 1997; de Almeida et al., 2001) to exogenously activate 5-HT1 receptors during optic nerve regeneration and measured its effects on regenerating RGC axons. Both the *htr1b* and *htr1d* receptors are expressed in RGC neurons prior to optic nerve transection and during regeneration at 24 hpt (Figures S2A-F and see mRNA expression in isl2b:GFP neurons, yellow dashed lines in Figures S2C’ and S2F’). Following optic nerve transection, we exposed larvae to zolmitriptan between 24 - 48 hpt to induce 5-HT1 receptor activation and assessed regeneration at 48 hpt. Compared to DMSO-treated control animals, treatment with 5 uM zolmitriptan significantly increased the number of optic nerves that re-innervated the contralateral tectum (Figures 5A-D, 1.6-fold change increase, compare DMSO controls with 5 uM zolmitriptan treated larvae) without inducing significant ectopic regrowth (Figure 5E), demonstrating that exogenous activation of 5-HT1 receptors enhances RGC axonal regeneration. However, increasing the agonist concentration to 50 uM caused a 2.3-fold increase of optic nerves that exhibited significant ectopic axonal regrowth (Figure 5E, compare DMSO controls with 50 uM zolmitriptan treated larvae). Specifically, in zolmitriptan treated larvae, regenerating RGC axon fascicles grew misguided from their pre-transection trajectory soon after exiting the injury site (Figures 5B-C, yellow arrowheads). In addition, in a small fraction (11%) of zolmitriptan treated animals, regenerating RGC axons initially extended towards the optic chiasm but then changed their course and continued on ectopic routes, ultimately failing to innervate the contralateral tectum (Figure 5B, white arrowhead). Lastly, exposure of larvae to 50 uM zolmitriptan did not affect the capacity of most regenerating axons to cross the optic chiasm and innervate the contralateral tectum (Figure 5D, no fold-change, compare DMSO controls with 50 uM zolmitriptan treated larvae). Thus, treatment with 5-HT1 receptor agonists at low concentrations (5 uM) enhances tectal re-innervation without causing significant ectopic regrowth, while high concentrations (50 uM) lead to severe anterior-posterior axonal misguidance. Combined, our results agree with the idea that in vivo 5-HT1 receptors can direct regenerating optic nerve axons, consistent with an instructive role. Finally, forced activation of 5-HT1 receptors at 32 hpt failed to elicit significant defects in optic nerve regeneration, including ectopic regrowth (Figures S2G-H), further underscoring the notion that serotonin signaling modulates RGC axon regeneration early during the regeneration process when RGC axons navigate towards the optic chiasm. In summary, our findings support the idea that 5-HT receptor-dependent neuromodulation plays an instructive and tightly regulated role during the early regrowth of RGC axons towards the optic chiasm.

**Figure 5.**
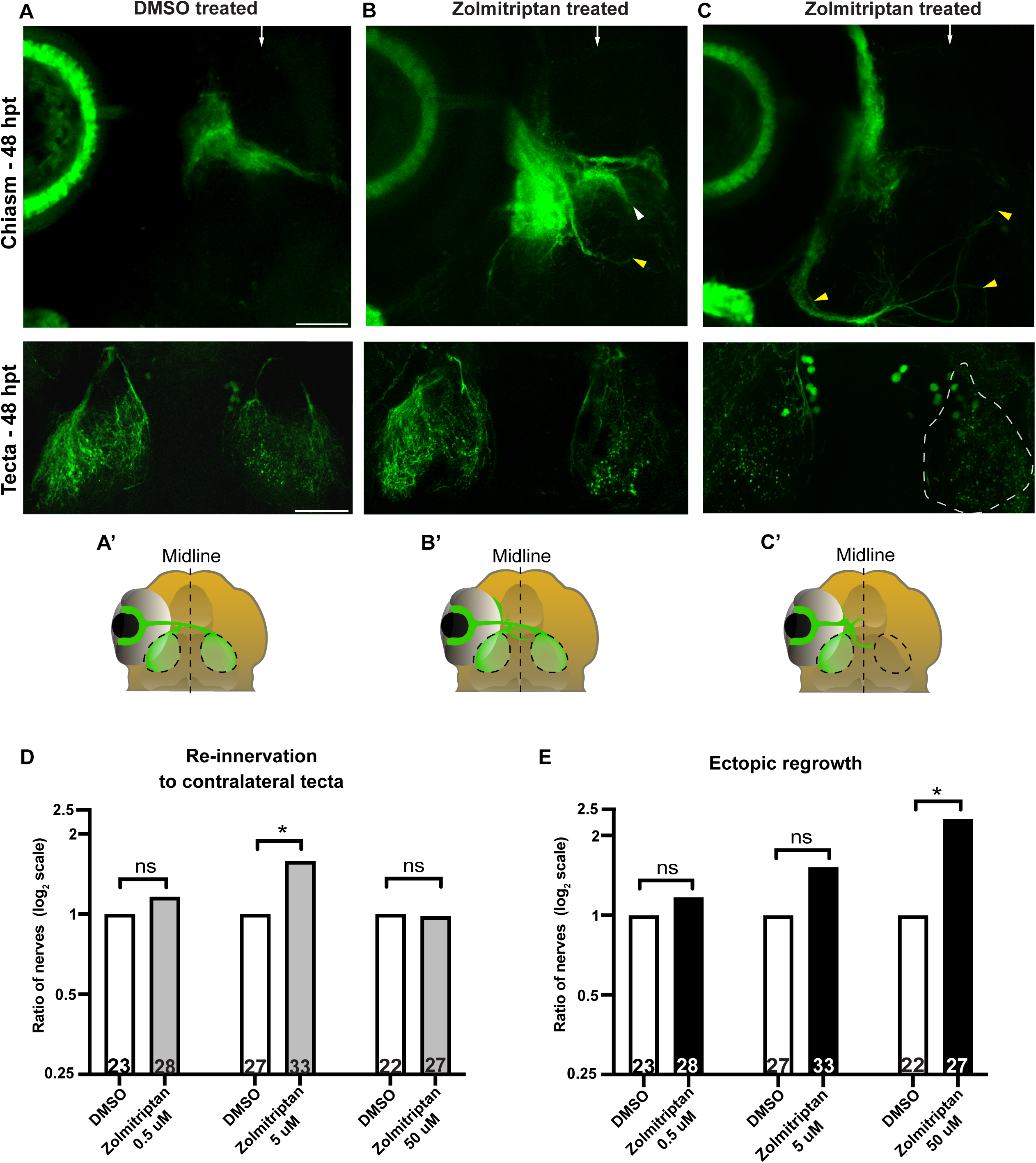
Exogenous activation of 5-HT1 receptors significantly increases axonal regrowth in a dose-dependent manner. (A - C) Fluorescent representative images and (A’ - C’) schematics of *Tg(Isl2b:GFP)* at 48 hpt of larvae exposed to (A) DMSO 0.3% and (B & C) 50 uM of agonist Zolmitriptan. Images in B and C are examples of two different types of regrowth observed when applying this small molecule. Scale bars = 50 uM. (B) Regenerating optic nerve of a larva treated with 50 uM Zolmitriptan shows axons exhibiting ectopic regrowth at the optic chiasm (white arrowhead). RGC axons also regrew misguided away from the chiasm, to a more posterior region of the brain (yellow arrowhead). Other RGC axons re-innervated both optic tecta by 48 hpt (bottom panel). (C) The regenerating optic nerve of a larva treated with 50 uM Zolmitriptan shows a significant amount of axon fascicles ectopically regrowing from the injury site and extending long distances away from the optic chiasm (yellow arrowheads). RGC axons did not re-innervate the contralateral tectum by the 48 hpt time point (bottom panel, dashed lines). White arrows depict the midline at the optic chiasm. (D) Quantification of optic nerve axonal re-innervation to contralateral tectum at 48 hpt in larvae treated at 24 hpt with DMSO 0.3% or the agonist Zolmitriptan. Bars represent the ratio of optic nerves that re-innervated the contralateral tectum in Zolmitriptan treated (gray bars) and control groups, plotted as a log2 scale. See Material and Methods for details on ratios and fold-change calculations. Data displayed in Figure 5D-E was obtained from larvae in the same treated groups. Statistics determined using the two-tailed Fisher’s exact test. Asterisks in figures represent statistical significance: * *P* < 0.05; ns, not significant. n=23 and n=28 for nerves treated with DMSO and 0.5 uM Zolmitriptan, respectively; n=27 and n=33 for nerves treated with DMSO and 5 uM Zolmitriptan, respectively; and n=22 and n=27 for nerves treated with DMSO and 50 uM Zolmitriptan, respectively. (E) Quantification of ectopic regrowth at 48 hpt in *Tg(Isl2b:GFP)* larvae treated at 24 hpt with DMSO 0.3% or Zolmitriptan. Bars represent the ratio of optic nerves showing ectopic regrowth in Zolmitriptan treated (black bars) and control groups, plotted as a log2 scale. See Material and Methods for details on ratios and fold-change calculations. Statistics determined using the two-tailed Fisher’s exact test. Asterisks in figures represent statistical significance: * *P* < 0.05; ns, not significant. n=23 and n=28 for nerves treated with DMSO and 0.5 uM Zolmitriptan, respectively; n=27 and n=33 for nerves treated with DMSO and 5 uM Zolmitriptan, respectively; and n=22 and n=27 for nerves treated with DMSO and 50 uM Zolmitriptan, respectively.

## Discussion

Here, we used a recently established optic nerve transection assay combined with the ease of small molecule treatment of larval zebrafish to uncover pathways that promote optic nerve regeneration. From a small molecule screen, we identified four small molecules that impaired optic nerve regeneration by selectively targeting 5-HT1 receptor signaling or other components of the serotonin signaling pathway. Previous studies in teleost fish have shown that after injury, serotonin receptor signaling can modulate neurite outgrowth of ex vivo retinal explants, as well as facilitate axonal regeneration of local spinal interneurons (Lima et al., 1994; Huang et al., 2021). To our knowledge, the findings reported here are the first to define an acute, in vivo role for serotonin receptor signaling in long-distance CNS axon and optic nerve regeneration.

Specifically, we find that 5-HT1 receptors promote optic nerve regrowth during the early stages of regeneration after regenerating RGC axons emerge from the nerve stump and begin to extend towards the optic chiasm. Moreover, 5-HT1 receptor signaling is dispensable during optic nerve development and in a separate CNS regeneration process, suggesting that 5-HT receptors play a selective role in the modulation of optic nerve regeneration. Finally, we show that ectopic activation of 5-HT1 receptors caused RGC axons to regrow ectopically, suggesting that serotonin receptor-dependent neuromodulation directs regenerating RGC axons towards the optic tectum.

### 5-HT1 receptor signaling promotes RGC axon regeneration but is dispensable during RGC axon development

An ongoing discussion in the field is whether individual molecules or even entire pathways promoting regeneration simply recapitulate developmental mechanisms. Over the past years, examples of genes and signaling pathways playing distinct roles in axonal growth during CNS regeneration compared to development have emerged (Becker and Becker 2007; Kusik et al., 2010; Wyatt et al., 2010). Gene expression analyses performed after optic nerve injury in adult zebrafish reveal that in addition to the expected increase in the expression of RGC intrinsic genes linked to development, growth-promoting genes not associated with this developmental process were also up-regulated (Veldman et al., 2007; Saul et al., 2010).

Our *in situ* hybridization data reveal that 5-HT1 *htr1aa* and *htr1ab* receptors are expressed in RGCs during the early stages of optic nerve regeneration. Similarly, previous work has demonstrated that the same 5-HT1 receptors are expressed in the retina and optic tectum during the development of the optic nerve (Pei et al., 2016). Given the expression profile of these receptors, we inhibited 5-HT1 receptors during the specific time window when RGC axons began extending towards the chiasm either during regeneration or development. Our findings that 5-HT1 receptor signaling functions during early optic nerve regrowth rather than development support the hypothesis that optic nerve regeneration is not merely a recapitulation of developmental programs.

It is therefore tempting to speculate that this differential requirement simply reflects a higher level of genetic redundancy among 5-HT receptors during development compared to regeneration. Consistent with this, in vivo knock-out studies of different 5-HT receptor subtypes in vertebrates (Saudou et al., 1994; Heisler et al., 1998) have failed to detect significant brain defects or even mild phenotypes in axonal patterning during development (Trakhtenberg and Goldberg, 2012). Moreover, pathways required for retinal axonal growth and guidance in zebrafish, such as heparan sulfate proteoglycans, are known to regulate this process with a tremendous amount of redundancy through the use of compensatory mechanisms during development (Poulain and Yost, 2015). Another possibility is that a different set of 5-HT receptors might function during optic nerve development. In fact, there are over 20 serotonin receptors in the zebrafish genome, most of which have their expression and function yet to be explored. Therefore, a thorough combinatorial analysis of 5-HT receptors will be critical to determine whether 5-HT receptors play a redundant role in optic nerve development.

### 5-HT1 receptors selectively promote early regeneration of optic nerve axons toward the optic chiasm

Pioneering work on the regrowth of optic nerve axons after injury in spontaneous regeneration models has described different stages of axon regeneration needed for the functional restoration of visual projections (Bohn and Reier, 1985; Stuermer et al., 1992; Kaneda et al., 2008). These optic nerve regeneration stages first include that after injury, RGC axons form a regenerative growth cone from the optic nerve stump; next, they extend toward the optic chiasm, across the chiasm, and finally, they extend toward the optic tectum where they re- innervate the optic tectum (Becker and Becker, 2007; Diekmann et al., 2015b). These regeneration stages can be distinguished at the transcriptional level, as reflected by gene expression, including the up-regulation of different transcription factors and pathways in RGCs during each stage (Dhara et al., 2019).

Our pharmacological inhibition of 5-HT1 receptors at distinct regeneration stages demonstrates that 5-HT1 receptors modulate the early extension of optic nerve axons towards the chiasm but are likely dispensable after crossing the chiasm. Thus, our findings that serotonin receptor signaling is selectively required for a particular regeneration stage are significant because they suggest that, similar to development, different genetic programs might control different stages of the optic nerve regeneration trajectory. For example, the mTOR pathway is another signaling pathway that might regulate early optic nerve regeneration in larval zebrafish. In mammals, mTOR signaling plays a critical role in potentiating the extension of regenerating RGC axons toward the chiasm (Park et al., 2008), and in adult zebrafish, it is sharply up-regulated when regenerating RGC axons grow towards the chiasm. Moreover, blocking mTOR activity inhibits optic nerve regeneration in adult fish (Diekmann et al., 2015a), a similar phenotype to the one we observe after blocking 5-HT receptor signaling. 5-HT receptors have previously been shown to interact with components of the mTOR pathway to regulate other CNS processes (Meffre et al., 2012; Teng et al., 2019), raising the possibility that these pathways might act together to promote the early stages of optic nerve regeneration.

### Serotonin receptor-dependent neuromodulation plays an instructive role in optic nerve regeneration

Our results demonstrate that inhibition of 5-HT1 receptors leads to reduced regeneration of optic nerve axons, while ectopic activation of 5-HT1 receptors leads to a partially opposite phenotype, resulting in increased regeneration and, at higher agonist levels, to misguided regrowth. This suggests that 5-HT1 receptors are critical for optic nerve regeneration and might, in fact, instruct the regeneration of RGC axons as they emerge from the nerve stump and navigate toward the optic chiasm. How may serotonin receptor-dependent signaling instruct regenerating axons during optic nerve regeneration? Recent work has shown that serotonin acts as a guidance cue when presented to growth cones in neuronal cultures. Specifically, growth cone repulsion is regulated through the 5-HT1B receptor and its inhibition of cyclic AMP (cAMP) levels (Vicenzi et al., 2021). Given the expression of different 5-HT1 receptor genes in RGC neurons during regeneration (this study), a plausible mechanism is that serotonin acts as a guidance cue along the optic nerve tracts and that its binding to 5-HT1 receptors on RGC axons repulses axons away from the retina and towards the optic chiasm. These receptors could, in turn, modulate cAMP signaling, which is pivotal for promoting RGC axon regeneration (Rodger et al., 2005; Hellstrom and Harvey, 2014). Alternatively, and not mutually exclusive, 5- HT receptor signaling might promote optic nerve regrowth to the CNS midline indirectly through neuromodulation of yet to be defined guidance cues. This interpretation is supported by previous research elucidating the role of 5-HT1B/1D receptor signaling in mediating the response of thalamocortical axons to netrin (Bonnin et al., 2007) and 5-HT2 receptors directing the midline crossing of commissural axons by regulating the translation of ephrinB2 (Xing et al., 2015).

Our findings are consistent with a model in which tight regulation of 5-HT1 receptor signaling levels is needed to effectively direct regenerating axons toward the chiasm. In this model, a dial indicator can be used to envision the 5-HT1 signaling levels that lead to a particular optic nerve regeneration phenotype. A dial pointing to normal 5-HT1 receptor signaling levels would correlate with the levels required for axons to extend and maintain their pre-transection trajectory after injury. Decreased levels of 5-HT1 receptor signaling, the levels likely attained after blocking 5-HT1 receptors with the antagonist WAY-100635, impede optic nerve regrowth. On the other hand, an increase in 5-HT1 receptor signaling, similar to the levels attained after ectopic activation with 5 uM of the agonist zolmitriptan, leads to enhanced optic nerve regrowth, culminating in higher rates of innervation to the contralateral tectum. Finally, increasing 5-HT1 receptor signaling levels further with 50 uM of zolmitriptan causes significant ectopic optic nerve regrowth, possibly due to a 5-HT1 receptor-mediated increase in the repulsion of axons to serotonin. Future work is required to determine whether 5-HT receptor signaling also promotes early optic nerve regeneration after injury in mammalian systems.

## Material and Methods

### Fish Maintenance and Ethics Statement

All animal protocols were approved by the University of Pennsylvania Institutional Animal Care and Use Committee (IACUC). *Danio rerio* transgenic lines were maintained in the Tübigen or Tupfel long fin (TLF) genetic background and raised as described in (Mullins et al., 1994). *Tg(isl2b:GFP)* (Pittman et al., 2008) larvae were maintained in dishes with phenylthiourea (1x PTU, 0.2 mM in E3 medium) beginning at 4 hours post fertilization and incubated in the dark at 29 °C to decrease melanocyte pigmentation.

### Transection Assay and Small Molecule Screen

At five days post-fertilization, larvae were anesthetized using 0.0053% tricaine in a PTU/E3 solution and then mounted on a glass microscopy slide using 2.5% low-melt agarose (SeaPlaque, Lonza) in a PTU/E3 solution containing 0.016% tricaine. Transections were performed on an Olympus SZX16 fluorescent microscope, as detailed in (Harvey 2019), with some modifications. Briefly, the optic nerve of the left eye was transected using a sharpened tungsten needle (Fine Science Tools), and the right eye was removed using sharpened forceps. Fish were detached from the agar and allowed to recover in dishes containing 1x Ringer’s Solution for 1 hr.

Complete transection of the optic nerve was confirmed by examining transected larvae at 22-24 hrs post-transection (hpt). Larvae that displayed no axonal-GFP remnants from the injury site to the tectum were added to 48-well plates and treated with DMSO 0.3% or small molecules at this time point unless otherwise noted. Small molecules used in the screen were obtained from the University of Pennsylvania High-Throughput Screening Core (Selleckchem Bioactive: 2100 FDA-approved/FDA-like small molecules)(Lamire 2023). The known targets of small molecules in the drug library were classified based on their genetic pathways using the PANTHER Classification System (pantherdb.org)(Mi and Thomas, 2009). To determine the number of genetic pathways that form part of the drug library, we submitted a target list provided by Selleckchem to the ‘Gene List Analysis’ section in PANTHER. Targets not recognized by the program were manually examined using the PANTHER ‘Prowler’ Pathway section. Stock solutions (100x frozen stocks in DMSO) were initially diluted 1:100 in E3, obtaining a 10x solution. 30 uL of this solution was then added into the wells, yielding a 10 uM drug concentration in 0.3% DMSO. Three unique small molecules were added to ‘small molecule pools’ to determine their effect on optic nerve regeneration. Investigators were blinded to the identity of the small molecules in each drug pool. Small molecules that lead to lethality, alteration of body morphology, or a significant reduction in responsiveness to touch during the treatment period were excluded from further testing. All small molecules of drug pools that impaired regeneration were then tested individually to validate the phenotypes observed using the assay described above. Individual small molecules that impaired regeneration were re- tested using a different batch of the same small molecule and confirming they impaired optic nerve regeneration.

### Individual Small Molecule Treatment

For individual treatment with small molecules, a 2.5 mM stock of WAY-100635 (Sigma W108) was prepared by dissolving a new vial of 10 mg of WAY-100635 powder in 7.42 ml of 100% DMSO. The stock solution was then further dissolved in PTU/E3 to a final concentration of 5 uM or 50 uM (0.3% DMSO final concentration). The stock solution of WAY-100635 was freeze-thawed a maximum of two times and then disposed of. A 50 mM stock of Zolmitriptan (Sigma SML0248) was prepared by dissolving a new vial of 10 mg of zolmitriptan powder in 696 uL of 100% DMSO. The stock solution was then further dissolved in PTU/E3 to a final concentration of 0.5 uM, 5 uM or 50 uM (0.3% DMSO final concentration). Zolmitriptan was applied to a 48- well plate dish containing 6 dpf larvae at different time points, as indicated in the results section. The stock solution of zolmitriptan was freeze-thawed a maximum of two times and then disposed of. Compounds were applied to wells in a 48-well plate dish containing fully transected 6 dpf larvae at different time points, as indicated in the results section. Control larvae received a solution with 0.3% DMSO in PTU/E3. Larvae were pooled and randomly assigned to control and experimental groups.

### Quantification of Optic Nerve (ON) Regrowth Phenotypes

For quantification of optic nerve re-innervation to the contralateral tectum following optic nerve transection, a defect was scored when a regenerating optic nerve (including any fascicle or individual axon part of this nerve) failed to re-innervate the optic tectum at 48 hpt. For quantification of optic nerves showing a misguided growth defect, an ectopic regrowth phenotype was scored when axons regrew at a 30 ° angle or more away from the stereotypical trajectory that regenerating axons follow toward the optic chiasm in pre-transected animals. Results are displayed in bar graphs representing the ratio of optic nerves that displayed a defect in drug treated or control treated groups. Ratios were calculated to normalize the results obtained from each experiment. The ratio for control treated groups (white bars) is always 1. The ratios for drug treated groups were calculated as follows:

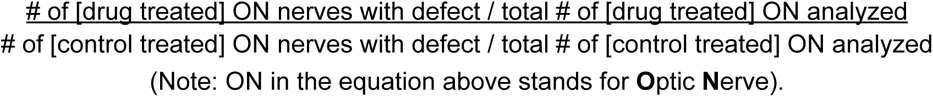

Quantification of mauthner cell axon regrowth at 96 hpt was performed as described in (Bremer 2019). The length of individual mauthner axons was measured in spinal cord hemisegments and compared between the control group (DMSO 0.3% treated) and drug treated group.

### Statistical Analysis

For statistical calculations of optic nerve regeneration phenotypes, we compared control and experimental groups on a contingency table and applied a two-tailed Fisher’s exact test for categorical outcomes between two samples using the online calculator provided at http://www.quantitativeskills.com/sisa/index.htm (Uitenbroek 1997). For statistical analysis related to mauthner axonal regeneration, we applied the One-way ANOVA test. Computations were performed using the Prism 10 software package from GraphPad. Graphs were generated using Prism 10 (GraphPad).

### Immunostaining and Confocal Imaging

Zebrafish larvae were stained as previously described (Harvey et al., 2023). Larvae were incubated with the primary antibody mouse anti-GFP (JL-8, 1:200, BD Biosciences) and secondary antibody goat anti-mouse Alexa 488 (1:500, Molecular Probes) overnight at 4 °C. Antibodies were diluted in 1% BSA and 1% DMSO in PBT. Stained larvae were mounted in Vectashield (Vector Laboratories) for confocal imaging. The brains of zebrafish larvae were imaged on a Zeiss LSM 880 confocal microscope using the 20x and 40x objectives. Image stacks of optic chasms or optic tecta were compressed into maximum-intensity projections. All images shown were adjusted for brightness and contrast, and color was assigned using Fiji.

### Fluorescent *in situ* Hybridization with Hybridization Chain Reaction (HCR)

*htr1aa, htr1ab, htr1b,* and, *htr1d* mRNA expression were detected using fluorescent *in situ* hybridization with HCR (Molecular Instruments, Los Angeles, CA, USA)(Choi et al., 2018). HCR probes, buffers, and hairpins were purchased from Molecular Instruments. Tg(isl2b:GFP) larvae were fixed at 5 dpf with 4% paraformaldehyde in PBS overnight at 4 °C in a rotating wheel. The staining was performed as previously described (Shainer et al., 2023) using B1 and B2 amplifiers with Alexa Fluor 546.

## Acknowledgments

We would like to tank Dr. Stout of the University of Pennsylvania CDB Microscopy Core for providing technical assistance, and Granato lab members for their comments and discussions.

## Author Contributions

Conceptualization: K.S.S., M.G.; Methodology: K.S.S., M.G.; Validation: K.S.S.; Formal analysis: K.S.S., M.G.; Investigation: K.S.S., M.B., J.M.; Resources: M.G.; Data curation: K.S.S., M.B.; Writing - original draft: K.S.S.; Writing - review & editing: K.S.S., M.G.; Visualization: K.S.S., J.M.; M.G.; Supervision: M.G.; Project administration: M.G.; Funding acquisition: M.G.

## Data Availability

All relevant data are within the article and its supplementary information files.

