## Supplemental Data for "Serotonin neuromodulation directs optic nerve regeneration"

A

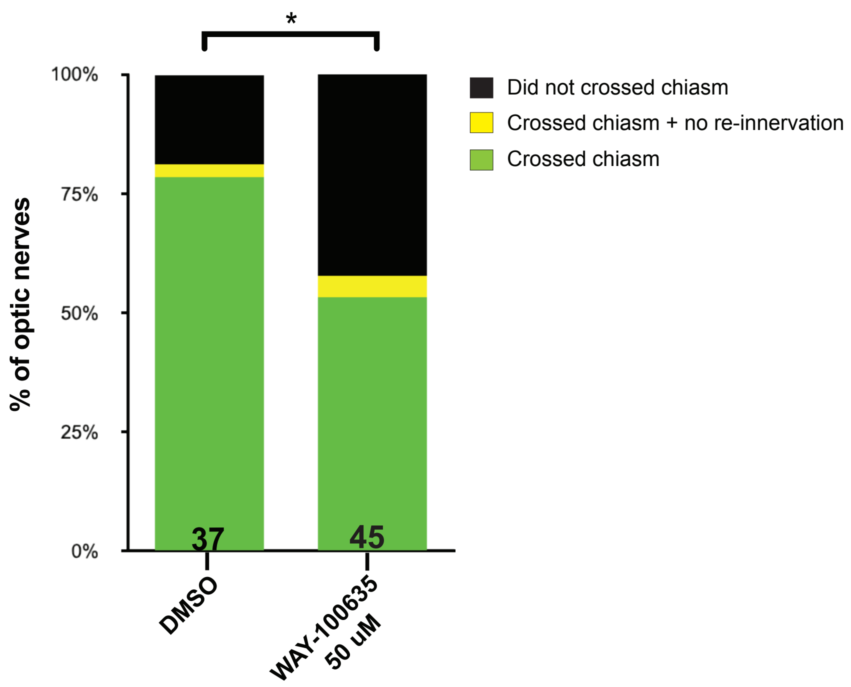


**Figure S1. Blocking 5-HT1 receptors significantly inhibited the regrowth of optic nerves to the optic chiasm.**

(A) Stacked bar graphs showing the percentage of optic nerves that crossed the midline by 48 hpt in *Tg(Isl2b:GFP)* larvae treated at 24 hpt with DMSO 0.3% or 50 uM of the antagonist WAY-100635. Green bars show the percentage of optic nerves that crossed the chiasm and re-innervated the contralateral tectum. Black bars show the percentage of optic nerves that did not cross the chiasm and did not re-innervated the contralateral tectum. Yellow bars show the percentage of optic nerves that crossed the chiasm but did not re-innervated the contralateral tectum. Data displayed was obtained from the same experimental groups analyzed in Figure 2D. n=37 and n=45 for nerves treated with DMSO and 50 uM WAY-100635, respectively. Statistics determined using the two-tailed Fisher's exact test. Asterisk in figures represent statistical significance: * $P < 0.05$; ns, not significant.

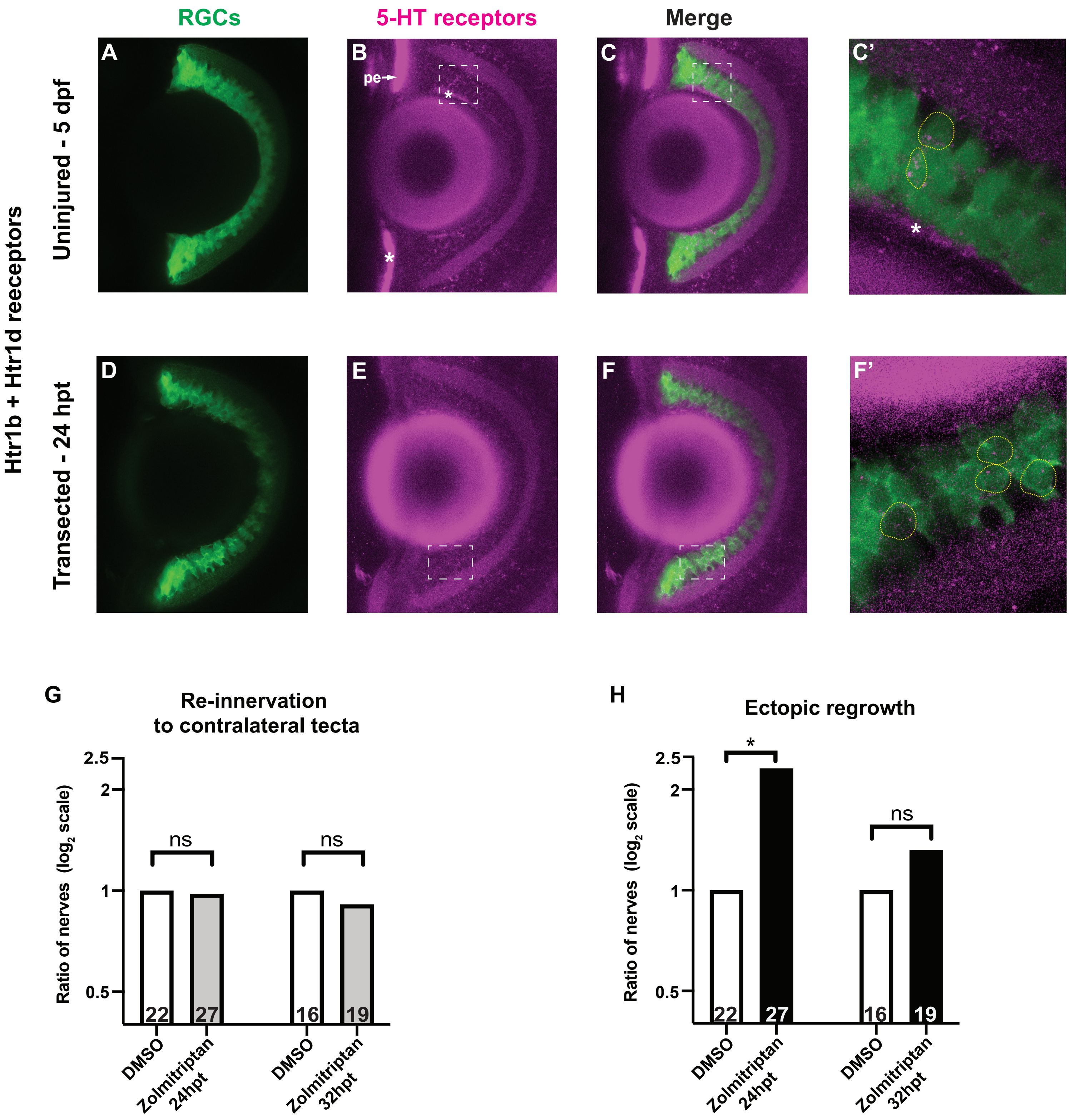


Saied-Santiago *et al.*, Supplemental Figure 2

**Figure S2. 5-HT1B and 5-HT1D receptor genes are expressed in RGCs at pre-transection and during optic nerve regeneration.**

(A - F) Representative images of retinas from *Tg(isl2b:GFP)* 5 dpf uninjured larvae (A-C', n=8) or larvae with transected optic nerves (D-F', n=3) at 24 hpt stained with an *in situ* hybridization HCR probe mix for *htr1b* and *htr1d* (magenta). Images shown are a maximum Z-projection of 10 optical sections (A-C) and 6 optical sections (D-F)(32 um), 40X. Dashed white boxes in (Figure B-C) is the area enlarged 1.5X in (C'), a merged maximum Z-projection of 15 optical sections (0.1 um). In (C'), yellow dashed lines outline cell bodies with mRNA expression. The brightness of the green channel was adjusted for better visualization of 5-HT1 receptor genes inside RGC neurons. White asterisks depict unspecific staining. The retinal pigmented epithelium (pe) was nonspecifically labeled by the amplifier with Alexa Fluor 546.

Dashed white boxes in (Figures E-F) is the area enlarged 1.5X in (F'), a merged maximum Z-projection of 6 optical sections (0.1 um). In (F'), yellow dashed lines outline cell bodies with mRNA expression. The brightness of the green channel was adjusted for better visualization of 5-HT1 receptor genes inside RGC neurons.

(G - H) Quantification of optic nerve regeneration phenotypes at 48 hpt in *Tg(isl2b:GFP)* larvae treated with DMSO 0.3% or 50 uM of the agonist Zolmitriptan at 24 and 32 hpt. Bar graphs and ratios observed were calculated as detailed in Figure 5D and in Material and Methods. Optic nerve regrowth phenotypes analyzed include optic nerve re-innervation to contralateral tecta (G, gray bars in Zolmitriptan treated) and ectopic axonal regrowth (H, black bars in Zolmitriptan treated). Data showing 50 uM Zolmitriptan at 24 hpt are identical to Figures 5D-E and shown here for visual comparison only. Statistics determined using the two-tailed Fisher's exact test. Asterisks in figures represent statistical significance: * $P < 0.05$; ns, not significant. n=22 and n=27, for nerves treated with DMSO and Zolmitriptan at 24 hpt, respectively. n=16 and n=19, for nerves treated with DMSO and Zolmitriptan at 32 hpt, respectively.
